# Cannabinoids is a “No-Go” While a Cancer Patient is on Immunotherapy; but is It Safe to Use Psychedelics During Cancer Immunotherapy?

**DOI:** 10.1101/2021.02.01.429102

**Authors:** E. Amit Romach, M. Nachliely, O. Moran, M. Brami, I. Lamensdorf

## Abstract

The use of Psychedelics by patients with cancer to relieve anxiety and depression has increased in the past few years. Since Psychedelics have immunomodulatory effects, their consumption among cancer patients should be carefully considered due to their potential negative effects on the tumor immune stroma, especially in view of the increase in the utilization of therapeutic approaches that are based on immune activation such as treatment with immune checkpoint inhibitors (ICIs). Preclinical data provided in this report indicate a potentially negative impact on tumor growth as a result of Psychedelics consumption during treatment with ICIs. Furthermore, our research suggests that the use of psychedelic agents (Lysergic acid diethylamide [LSD] or Psylocibin) might diminish the beneficial therapeutic benefits of ICIs.

It might be necessary to expend this line of research in order to validate these findings, in view of the increase use of cannabinoids and psychedelics among cancer patients, some of them being treated with immune-based modalities.

## Introduction

Immunomodulation is a major pathway used by cancer cells to silence the immune system’s natural response and to continue cancer cell proliferation. The field of immunotherapy in general, and checkpoint inhibitors in particular, has progressed significantly during the past several years and is perceived as a new promise for cancer patients. It includes various therapeutic mechanisms that harness the immune system in order to control malignancy. One major class of immunotherapy drugs, that has been approved for clinical use, are antibodies against the Programmed Death receptor 1 (PD1) or its ligand (PD-L1), which directly inhibit the PD1/PD-L1 interaction. Another class of approved immunotherapy is neutralizing antibodies targeting the immune checkpoints T-lymphocyte-associated protein 4 (CTLA-4). In subset of responding patients, this inhibition resulted in activation of the immune system, and as a result diminution of the proliferative process [1,2].

Psychedelics and Cannabinoids are being used extensively by cancer patient to relieve stress, address cachexia, pain and inflammation. Although not regulatory approved or clinically validated, therapeutic antineoplastic effect has been attributed to cannabinoids, potentially as an auxiliary to chemotherapeutic agents. However, accumulating preclinical and clinical findings clearly demonstrated that their use in patients receiving immune-based therapies (ICIs) should be cautiously considered, in view of increasing evidence that cannabinoids diminish ICIs activity.

There is evidence that psychedelic drugs reduce anxiety and depressive symptoms, leading to a decrease in cancer-related demoralization and hopelessness, improved spiritual well-being, and increased quality of life [3]. However, currently, information about their in vivo effect on cancer growth and the potential interaction between Psychedelics and ICIs is lacking.

Psychedelics (serotonergic hallucinogens) are powerful psychoactive substances that alter perception and mood and affect numerous cognitive processes. They are generally considered physiologically safe and do not lead to dependence or addiction. Depending on the specific drug, it might affect non-specifically a large spectrum of receptors. However, it is well established, that their main effect is modulated via activation of the serotonin 5-HT2A receptor.

5-HT2A receptor is known to be involved in cell proliferation and it is known that changes in serotonergic innervation may modulate the immune response to inflammatory as well as cancerous processes [4,5].

Classical psychedelics such as LSD and DMT (N,N-dimethyltryptamine) have been shown, in vitro, to exert anti-cancer and anti-inflammatory effects through the modulation of innate and adaptive immune processes [6–8].

In view of the increase in their clinical utilization and the concern as potential pro-proliferative effect modulated via the serotonergic system and their potential immune suppressive activity, we have conducted a small scale evaluation utilizing two psychedelic compound/extracts: Lysergic acid diethylamide (LSD) and “magic mushroom” extract containing Psylocibin in vitro and in vivo utilizing pre-clinical models, with or without PD1 inhibitor, in order to investigate whether there is a merit to the above-mentioned concerns.

## Methods

### Cytotoxicity assay

Cytotoxicity of a psychoactive plant extract on breast cancer cell line was evaluated.

E0771 cells were plated in a 96 well plate, in their culture medium. Cells were allowed to attach for 16-24 hours at 37°C and 5% CO_2_. Thereafter, culture medium was discarded, and fresh assay medium was added to the cells, supplemented with the Test Item (103 μg/ml Psylocibin or 6.8 μg/ml LSD)

Cells were incubated for another 48 hours at 37°C and 5% CO_2_. Finally, fresh culture medium was added to the cells along with 50 μL XTT reagent.

The Optical Density (OD) was measured using a plate reader once Vehicle treated cells reached the OD range of 0.5-1.5 OD at 450 nm wavelength.

### Syngeneic Mouse Model

The anticancer activity of psychoactive plant extract and LSD was evaluated in a syngeneic model of E0771 breast cancer in mice, alone or in combination with anti-PD1.

Mice were treated PO with a mushroom extract containing Psilocybin (0.685 μg/mouse) and LSD (10.3 μg/mouse) 5 days a week and injected IP with Anti PD1 twice a week.

## Results

Both Psilocybin (Fig. 1) and LSD (Fig. 2) induced cell death in E0771 breast cancer cell line in vitro.

However, Psilocybin and LSD, contrary to the observed in vitro effect, induced tumor growth in a syngeneic model of E0771 breast cancer in mice in vivo (Fig. 3).

**Fig. 1:**
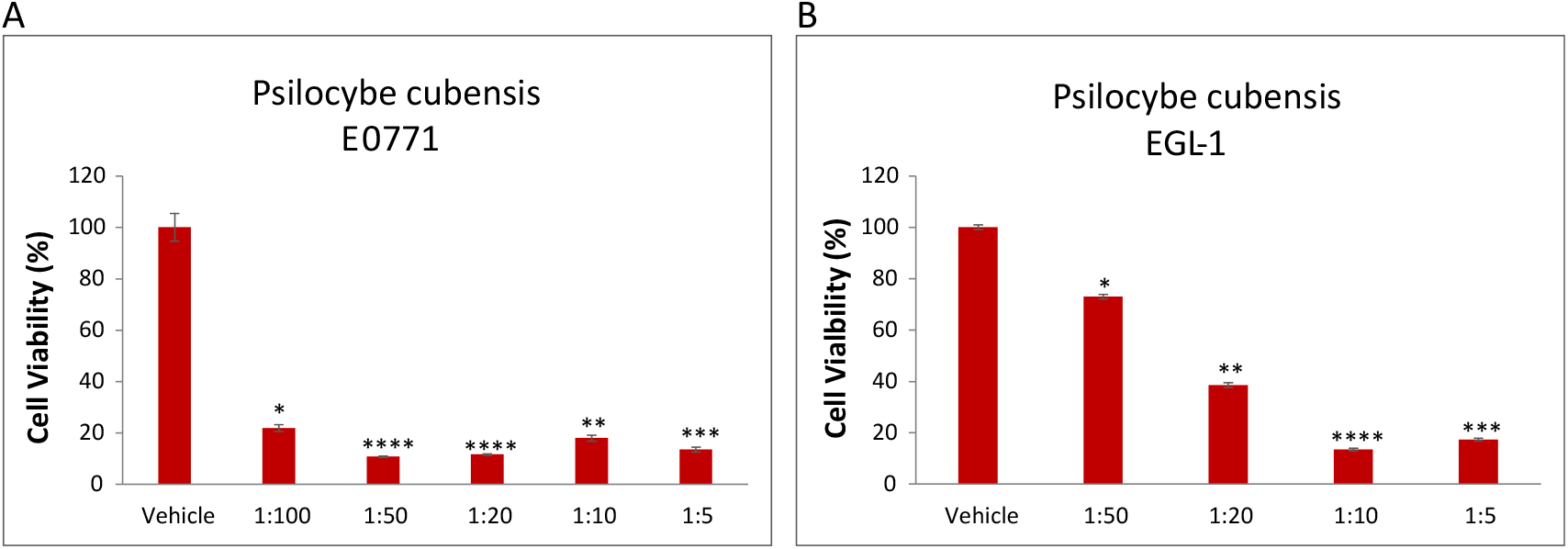
Psilocybe cubensis (Psilocybin) reduced the viability of E0771 breast cancer cells (A) and EGL-1 cells (B) in vitro.

**Fig. 2:**
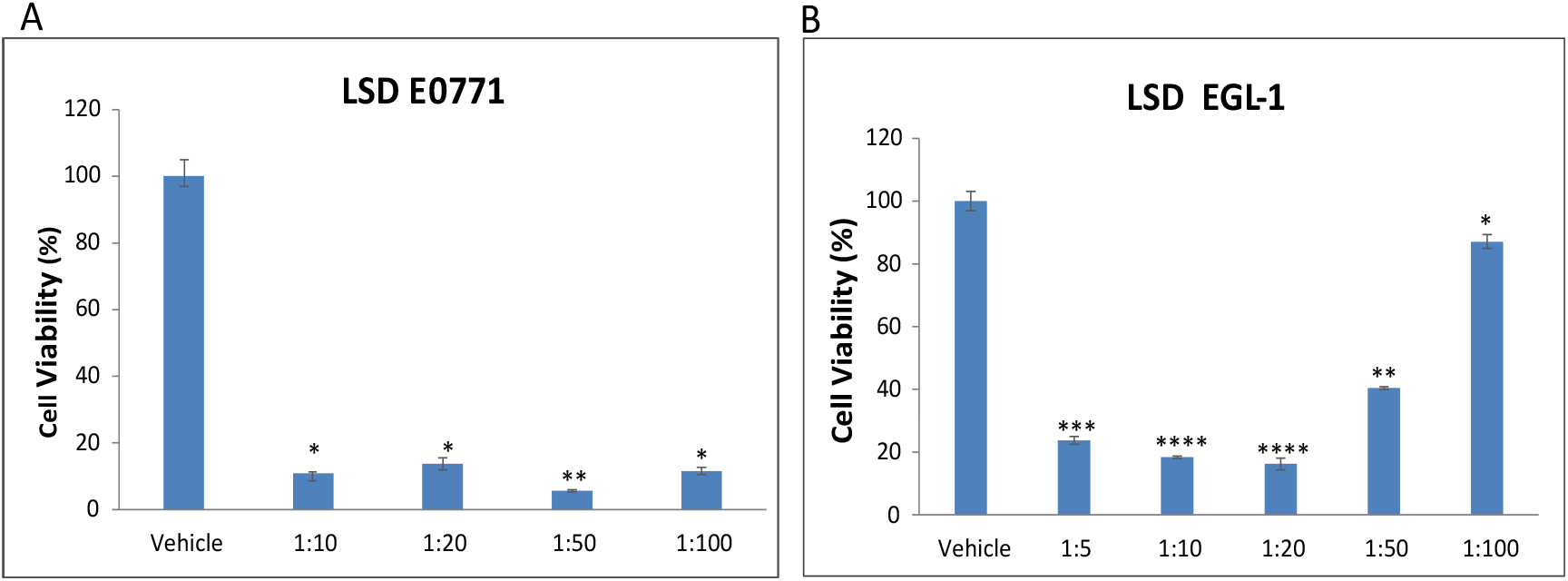
LSD (Lysergic acid diethylamide) reduced the viability of E0771 breast cancer cells (A) and EGL-1 cells (B) in vitro.

**Fig. 3:**
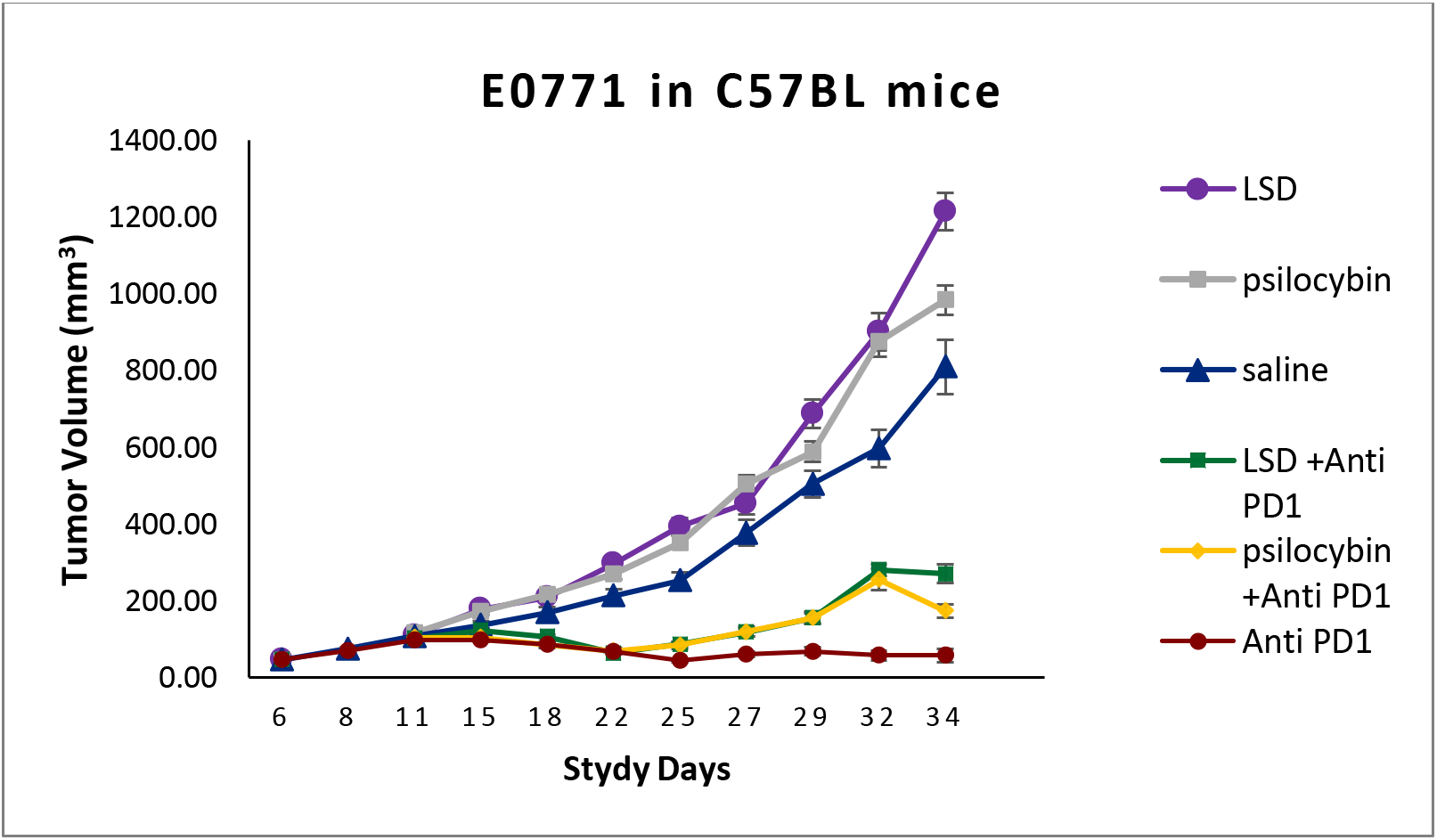
Psilocybin and LSD induce tumor growth and the combination of Psilocybin /LSD with Anti PD1 interfere with Anti PD1 activity in E0771 mice model.

## Conclusions

Psychedelics are used by patients with cancer to help relieve cancer symptoms and treatment-related side effects. However, Psychedelics immunomodulatory properties described in this report, suggest Psychedelics could conflict with cancer treatment in general and specifically with immunotherapy.

This information can be critical for a large group of patients and requires caution when starting immunotherapy and considering utilization of psychedelic agents even when ICIs are not being used.

